# Validating an Immunoassay to Measure Fecal Glucocorticoid Metabolites in Yellow-Bellied Marmots

**DOI:** 10.1101/2024.05.20.595012

**Authors:** Xochitl Ortiz-Ross, Hash Brown Taha, Emily Press, Sarah Rhone, Daniel T. Blumstein

## Abstract

The yellow-bellied marmot (*Marmota flaviventer*) study at the Rocky Mountain Biological Laboratory near Crested Butte, Colorado, USA is the world’s second longest study of free-living mammals. Quantifying physiological stress is essential for understanding their health, reproductive success, and survival in a variable environment. Historically, we used a validated radioimmunoassay (RIA) to measure fecal glucocorticoid metabolites (FGMs). Given the costs and risks of working with radioisotopes, we have shifted to a more sustainable method. Here we evaluate the suitability of two competitive enzyme-linked immunosorbent assays (ELISA) from Cayman Chemical Company (CCC) and Arbor Assays (AA) to measure corticosterone levels in FGMs. The findings revealed that the AA ELISA, unlike the CCC ELISA, consistently matched the RIA in terms of accuracy across high and low corticosterone concentrations, demonstrated superior assay parameters, showed the highest correlations with RIA results and effectively captured the annual variations in FGM concentrations, indicative of its reliability for use in longitudinal studies. We further analytically validated the usage of the AA ELISA for FGMs, confirming its efficacy without matrix effects, thus establishing its suitability for ongoing and future studies of FGMs in marmots. The transition to the AA ELISA from the RIA ensures continued data integrity while enhancing safety and environmental sustainability.

## 1. INTRODUCTION

Quantifying physiological stress in wild animals is crucial for several reasons. It serves as a gauge for the overall health and resilience of individual animals. Monitoring stress levels can help us understand how environmental pressures such as habitat degradation, pollution, or predation impact the population’s viability. Furthermore, physiological stress can serve as a key indicator of ecological disturbance, allowing us to assess the stability and functionality of ecosystems (Boonstra, 2012; Karaer et al., 2023; Romero and Wingfield, 2015; Sapolsky et al., 2000; Sheriff et al., 2011). Therefore, a comprehensive understanding of physiological stress in wild animals can better inform conservation efforts and ecosystem management strategies, ultimately contributing to the preservation of biodiversity. Such data becomes even more useful when it is collected over long periods of time to assess trends.

The population of yellow-bellied marmots (*Marmota flaviventer*) around the Rocky Mountain Biological Laboratory near Crested Butte, Colorado, USA is the world’s second longest, continually studied mammal (Armitage, 2014; Blumstein et al., 2013). This large ground-dwelling squirrel is notable for their obligatory hibernation and susceptibility to significant seasonal variations in food resources, predation, and weather. Behavioral and morphological data has been collected on uniquely identified yellow-bellied marmot individuals since 1962. In 2002, we added the regular collection of physiological and genetic data, including fecal samples from which we extract fecal glucocorticoid metabolites (FGMs). For the past 20 years we have been analyzing marmot FGM levels using a corticosterone radioimmunoassay (RIA; MP Biomedicals) following (Smith et al., 2012). We were forced to halt analysis using RIAs and decided to turn to enzyme-linked immunoassays (ELISAs). Given the longitudinal nature of our data, it is important that this new data source is comparable to the previous analyses to maintain data continuity.

RIA and ELISA are both immunoassays used for measuring a biomarker of interest, often in biological samples. RIA, one of the first immunoassay techniques developed, uses radioactively labeled antigens to detect the presence of antibodies in a sample. The degree of radioactivity allows for quantification of the antibody concentration. By contrast, ELISA uses an enzyme-linked antigen or antibody as a marker instead of a radioactive isotope. The enzyme, in the presence of its substrate, produces a measurable product, typically a color change, that correlates with the substance concentration in the sample. Both methods rely on the specific binding between antigen and antibody but differ in their detection strategies—RIA using radioactivity and ELISA using a colorimetric change.

In general, RIAs are currently less readily available, present greater safety and sustainability concerns, and strict radioactive handling regimens, rendering them less convenient compared to ELISAs. On the other hand, ELISAs have become more popular due to their non-radioactive nature, making them safer and easier to handle. Furthermore, ELISAs can be performed in different formats such as direct, indirect, sandwich, or competitive, each suited for different biomarkers. However, both types of immunoassays require precision and careful handling to ensure their accuracy (Cox et al., 2004; Porstmann and Kiessig, 1992).

While several studies have made a similar switch from RIA to ELISA (Al-Dujaili et al., 2009; Elder et al., 1987; Glucs et al., 2018; Kinn Rod et al., 2017), they do not usually involve such a long-term data set. This is important because extended longitudinal data provide a more comprehensive understanding of hormonal patterns and their fluctuations over time. It allows for the assessment of seasonality, reproductive cycles, and the effects of environmental stressors across the lifespan of the subjects, offering invaluable insights into the physiological resilience and health of the population under study. Furthermore, most ELISAs are validated and used for blood, while there is little research that has been published on the use of fecal samples. FGMs are particularly useful measures of glucocorticoids (GCs, e.g., corticosterone, cortisol) because feces are easy to obtain through minimally invasive and non-invasive sampling in animal populations (see methods). Lastly, no research has been published on validating ELISAs for measuring FGMs from yellow-bellied marmots.

Thus, to continue harnessing two decades of data while switching to safer, more convenient and sustainable methods, and in an effort to guarantee data accuracy, we compared two competitive ELISA kits from Cayman Chemical Company (CCC) and Arbor Assays (AA) with the RIA results we had previously obtained (using a mix of years from 2016 to 2020). More specifically, we conducted detailed comparison of FGM concentrations, a thorough examination of assay parameters, a robust correlation and covariate analysis, and a rigorous analytical validation of chosen ELISA to ensure the accuracy and reliability of long-term stress data in yellow-bellied marmots.

The AA ELISA emerged as the superior method in our comparative analysis, consistently matching the RIA’s accuracy across high and low FGM concentrations and proving to be the most reliable for detecting nuanced stress responses in yellow-bellied marmots. Its superior assay parameters and robust analytical validation outcomes showcasing a lack of a matrix effect, firmly establish it as the method of choice for our ongoing longitudinal studies. Long-term longitudinal data sets that were previously used to obtain data from RIAs may be able to switch to using the AA ELISA to validate their new data.

## 2. MATERIALS AND METHODS

### 2.1 Sample collection and FGM extraction

We collected fecal samples from wild yellow-bellied marmots in the vicinity of the Rocky Mountain Biological Laboratory (RMBL), located in the East River Valley, in Gunnison County, Colorado USA. Fecal samples have been collected since 2002. During the active season, spanning from May to August, we aimed to capture marmots on a bi-weekly basis using Tomahawk traps positioned at burrow entrances. Upon reaching each trap, the marmots were carefully transferred into canvas handling bags for weight measurement and sex determination, with their reproductive states meticulously recorded (females categorized as pregnant, lactating, or non-reproductive, while males were classified as scrotal or abdominal). At the time of initial capture, all marmots were uniquely marked using Nyanzol cattle dye and uniquely numbered ear tags (Armitage, 1982). Using a plastic Ziplock bag, fecal samples were collected from the trap (typically within 2 hours post-defecation) or, opportunistically, directly from the bagged marmot and immediately stored on wet ice until they were transported back to the lab and frozen at −20°C. Annually, during the month of August, the samples were transported on dry ice to the Blumstein Lab at the University of California (UCLA) for hormone extraction (as described in (Smith et al., 2012)). We developed our intial RIA in collaboration with researchers at the San Diego Zoo (Blumstein et al., 2006), but for most of the past 20 years RIAs were conducted by the Wasser lab at University of Washington.

### 2.2 Corticosterone assessment

We measured corticosterone from FGM extracts using a radioimmunoassay (RIA) from MP Biomedicals (MPB; Cat # 0712010-CF) or two different competitive ELISAs specific for corticosterone from CCC (Cat # 501320) and AA (Cat # K014-H1/H5). The RIA samples were diluted 4-fold, whereas CCC and AA samples were diluted 8-fold. All further immunoassay steps were conducted according to the manufacturer’s instructions. Data from the CCC immunoassay was analyzed according to the manufacturer’s protocol. Data from the AA immunoassay was analyzed using the following website https://myassays.com/arbor-assays-corticosterone-enzyme-immunoassay-kit-high-sensitivity.assay.

### 2.3 Assay parameters

We calculated the limit of blank (LoB), lower limit of detection (LLoD), lower limit of quantification (LLoQ) and upper limit of quantification (ULoQ) as described previously (Armbruster and Pry, 2008; Taha, 2024) and the intra- sand inter-assay coefficient of variations (CV) using the standard curve from four different runs. Total (TE) and relative (RE) error were calculated as described previously (Taha*-Dutta*-Hornung* et al., 2023; Kat and Els, 2012; Taha, 2024; Westgard et al., 1974).

### 2.4 Spike Recovery

To test spike recovery, fecal lysates were diluted ≤ the hypothetical lower limit of quantification (LLoQ) using the assay’s buffer to avoid exceeding the from signal highest standard calibrator (Smith et al., 2012), and each sample was spiked with 39.0 pg/mL (low spike), 157.0 pg/mL (medium-low spike), 595.0 (medium-high spike), and 1337.0 pg/mL (high spike) of the recombinant standard calibrator provided with the ELISA from AA. Percentage recovery was calculated using the below formula.

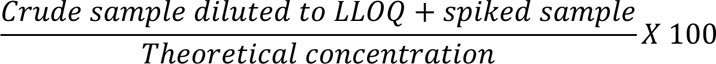

### 2.5 Dilution linearity

To test dilution linearity, undiluted fecal lysates were spiked to the ULoQ using the highest recombinant standard calibrator and diluted 2-, 4-, 8- and 16-folds in the assay’s buffer. Percentage recovery was calculated using the below formula.

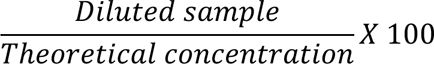

### 2.6 Parallelism

Two FGM extracts were diluted serially 2-, 4- and 8-folds using the assay’s buffer. Percentage recovery was calculated similarly to dilution linearity. Parallelism was observed by evaluating relative accuracy using linear regression curves from the diluted samples and standard calibrators.

### 2.7 Statistical analysis

We performed Pearson’s correlation tests to assess the associations among FGM concentrations obtained from the three immunoassays. Additionally, we considered year, sex, and age as covariates in our linear regression models to account for potential confounding factors. These models were also employed to determine the coefficient of determination (R²), which assesses the proportion of variance explained in the FGM concentrations, and to evaluate the parallelism between the standard calibrators and the diluted samples using the AA immunoassay. All analyses were conducted in RStudio (version 4.3.1, (Team, 2023)).

## 3. RESULTS

### 3.1 FGM levels

We first aimed to compare the levels of FGM using the three immunoassays. The quantification revealed that, although both the RIA and the CCC ELISA had an overall lower mean (mean ± SEM: 1601.7 ± 80.7 and 2017.1 ± 102.3 pg/mL, respectively) than the AA ELISA (mean ± SEM: 2971.3 ± 159.3 pg/mL). However, we observed that both the RIA and the AA ELISA detected similar values at the highest and lowest end of concentrations (**Figure 1**). This supports our experience with the CCC ELISA; in four different runs, and after testing multiple dilution factors, all samples (n = 48) still gave values above the highest standard, precluding our ability to quantify their concentrations. This suggests that the CCC ELISA may not be reliable for quantifying samples with high FGM concentrations or those derived from yellow-bellied marmots under stressful conditions.

**FIGURE 1.**
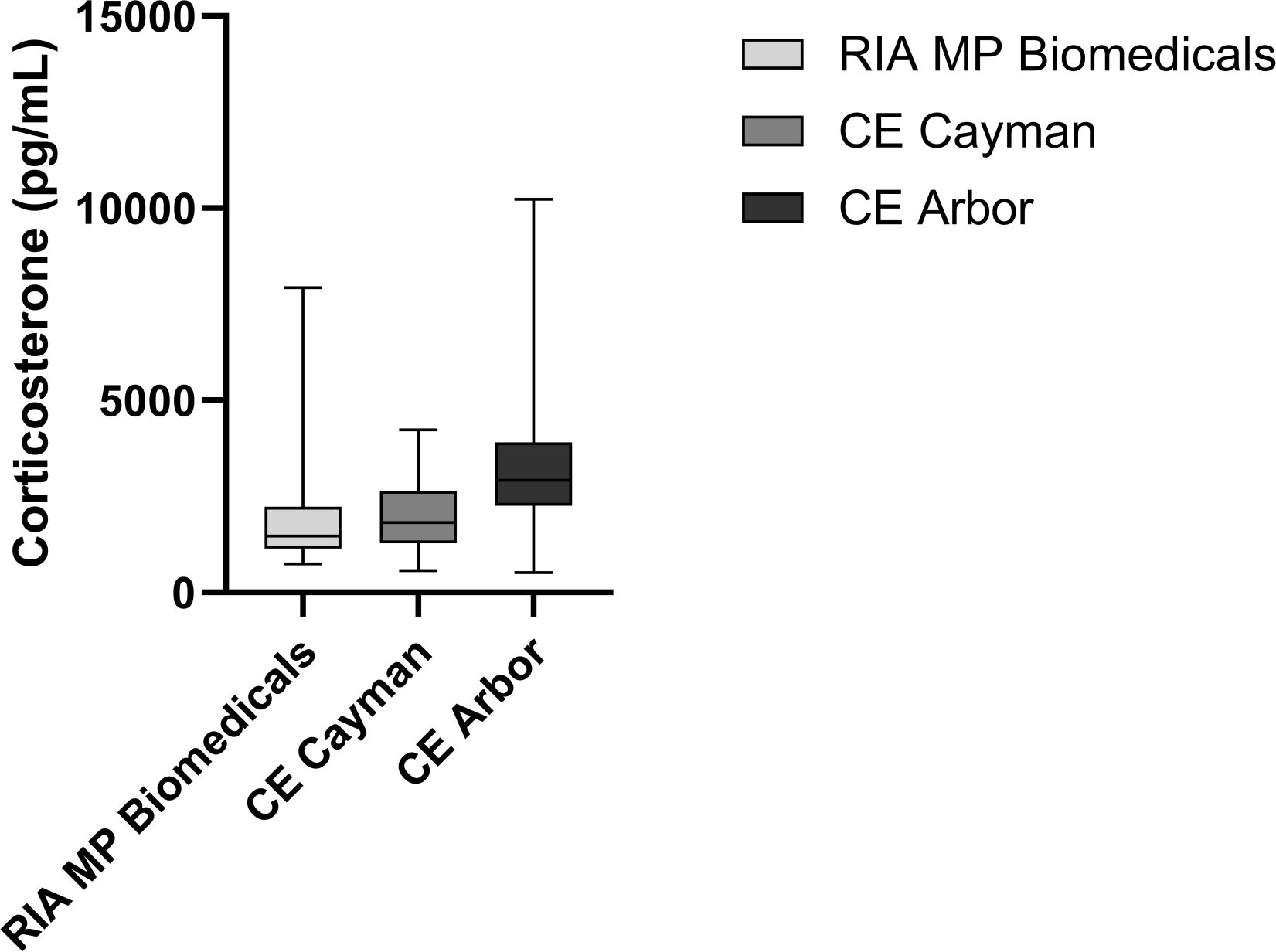
Evaluation of yellow-bellied marmots’ fecal glucocorticoid metabolites (FGM) using a radioimmunoassay (RIA) or two competitive enzyme-linked immunosorbent assays (ELISA) from Cayman Chemical Company or Arbor Assays specific for corticosterone. CE – Competitive ELISA.

### 3.2 Assay parameters

We next evaluated how each ELISAs performed across different analytical parameters, including the LoB, LLoD, LLoQ, ULoQ, and stability, using intra- and inter-assay CVs (**Table 1**), as well as TE and RE (**Tables 2-3**).

**Table 1.** Analytical parameters for detection of fecal glucocorticoid metabolites (FGM) using the two competitive immunoassays. The limit of blank (LoB), lower limit of detection (LLoD), lower limit of quantification (LLoQ) and upper limit of quantification (ULoQ) are quantified as previously described (Armbruster and Pry, 2008; Dutta et al., 2023). CV – coefficient of variation; ELISA – enzyme-linked immunosorbent assay.

| Assay | Company | LoB<br>(pg/mL) | LLoD<br>(pg/mL) | LLoQ<br>(pg/mL) | ULoQ<br>(pg/mL) | Intra-<br>assay<br>CV | Inter-assay<br>CV |
| --- | --- | --- | --- | --- | --- | --- | --- |
| Competitive<br>ELISA | Cayman<br>Chemical<br>Company | 37.1 | 191.1 | 825.2 | 5797 | 16.3% | 16.3% |
| Competitive<br>ELISA | Arbor<br>Assays | 22.1 | 27.9 | 70.9 | 5189 | 12.4% | 13.6% |

**Table 2.** Calculated concentrations, total error (TE) (Westgard et al., 1974), and relative error (Kat and Els, 2012) of fecal glucocorticoid metabolites (FGM) using the competitive enzyme-linked immunosorbent assay from Cayman’s Chemical Company.

| Corticosterone<br>(pg/mL) | Assay 1 |  | Assay 2 |  | Assay 3 |  | Assay 4 |  | TE<br>(%) | RE<br>(%) |
| --- | --- | --- | --- | --- | --- | --- | --- | --- | --- | --- |
| 5000 | 4735 | 7175 | 4093 | 6677 | 4361 | 9625 | 4423 | 8998 | 113.16 | 25.22 |
| 2000 | 2024 | 2052 | 1949 | 1797 | 1919 | 1689 | 2028 | 1780 | 8.71 | 4.76 |
| 800 | 776.6 | 691.9 | 727.2 | 928 | 750.1 | 857.4 | 796.7 | 766.5 | 17.12 | 1.65 |
| 320 | 321.7 | 314.8 | 356 | 308.3 | 386.6 | 260.9 | 317.4 | 279.9 | 24.19 | 0.56 |
| 128 | 142.3 | 129.2 | 134 | 124.6 | 150.7 | 117.2 | 160.9 | 145.6 | 30.49 | 7.86 |
| 51 | 48.56 | 51.91 | 45.72 | 42.74 | 49.8 | 50.15 | 52.3 | 43.92 | 8.55 | 5.61 |
| 20 | 22.37 | 15.97 | 24.01 | 26.73 | 21.66 | 16.25 | 23.94 | 10.68 | 54.74 | 1.01 |
| 8 | 10.94 | 6.183 | 5.222 | 8.344 | 8.61 | 8.779 | 8.2 | 7.2 | 42.90 | 0.82 |

TE (%) = ((calculated concentration – actual concentration) + 2 SD)/actual concentration) X 100;
RE (%) = ((calculated concentration – actual concentration)/actual concentration) X 100.

**Table 3.** Calculated concentrations, total error (TE) (Westgard et al., 1974), and relative error (Kat and Els, 2012) of fecal glucocorticoid metabolites (FGM) using the competitive enzyme-linked immunosorbent assay from Arbor Assays.

| Corticosterone<br>(pg/mL) | Assay 1 |  | Assay 2 |  | Assay 3 |  | Assay 4 |  | TE<br>(%) | RE<br>(%) |
| --- | --- | --- | --- | --- | --- | --- | --- | --- | --- | --- |
| 5000 | 4988 | 4710 | 5613 | 4653 | 4594 | 4734 | 4850 | 4796 | 10.34 | 2.66 |
| 2500 | 2650 | 2575 | 2541 | 2633 | 2655 | 2519 | 2749 | 2598 | 10.47 | 4.60 |
| 1250 | 1248 | 1248 | 1191 | 1166 | 1350 | 1271 | 1127 | 1251 | 9.57 | 1.48 |
| 625 | 593.7 | 615.5 | 570.5 | 729.8 | 611.4 | 622.1 | 590 | 697.6 | 18.39 | 0.61 |
| 312.5 | 326.9 | 309.6 | 338.2 | 276.4 | 317.8 | 298.2 | 271.8 | 357.8 | 18.79 | 0.13 |
| 156.25 | 165.3 | 151.2 | 138.9 | 199.9 | 152.1 | 139 | 138.2 | 169.9 | 27.46 | 0.36 |
| 78.1 | 87.91 | 73.18 | 52.76 | 89.08 | 86.11 | 99.04 | 68.71 | 88.95 | 41.30 | 3.35 |
| 39.05 | 41.13 | 27.03 | 41.53 | 32.04 | 31.94 | 35.94 | 25.6 | 58.87 | 48.75 | 5.86 |
| 19.5 | 33.02 | 12.25 | 22.89 | 23.49 | 21.74 | 18.53 | 18.05 | 19.3 | 69.66 | 8.51 |

TE (%) = ((calculated concentration – actual concentration) + 2 SD)/actual concentration) X 100;
RE (%) = ((calculated concentration – actual concentration)/actual concentration) X 100.

In our hands, the CCC ELISA exhibited a large LoB, LLoD, and LLoQ due to variability in the zero and the lowest standard calibrators across different runs, as well as higher CVs, TEs and REs. However, the AA ELISA resulted better overall metrics (lower LoB, LLoD, LLoQ, CVs, TEs and REs), suggesting that this immunoassay is more reliable for our purposes.

### 3.3 Correlational analysis

Previously, we successfully validated the use of the RIA for the quantification of FGMs extracted from feces of yellow-bellied marmots. This validation was substantiated by observing a marked increase in FGM concentration in response to both biological and physiological stressors, as well as after trapping events (Smith et al., 2012). Such findings indicate that the RIA method is sensitive and specific enough to detect genuine elevations in stress levels within yellow-bellied marmots. However, due to constraints preventing the continued application of RIA, we sought to identify which ELISA has comparable accuracy by evaluating their association with the RIA assay.

We observed statistically significant moderate to strong correlations among the three immunoassays (**Figure 2A-C**). However, the AA ELISA showed the highest correlation with the RIA (**Figure 2A**), suggesting it is the best choice for replacing the RIA for quantifying physiological stress levels in yellow-bellied marmots’ FGMs.

**FIGURE 2.**
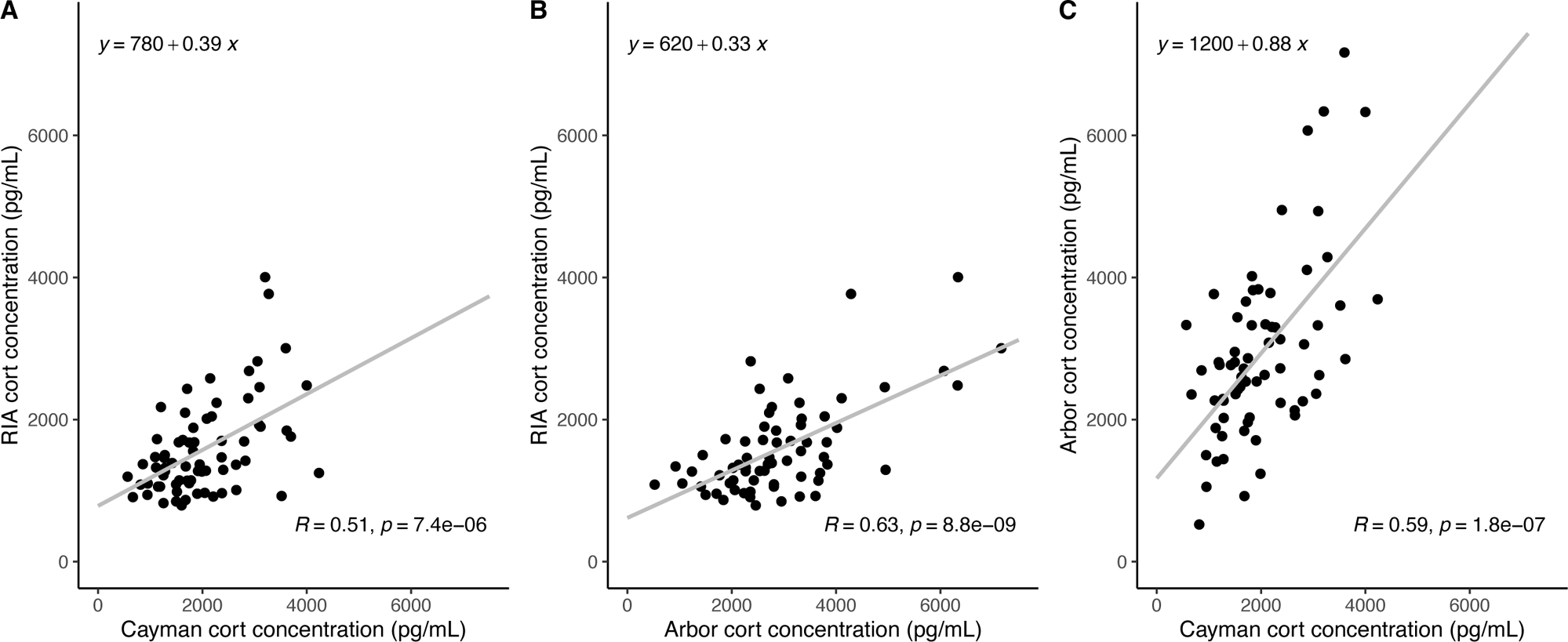
Association of corticosterone levels in yellow-bellied marmots’ fecal glucocorticoid metabolites (FGMs) among the three included immunoassays. Cort – corticosterone; RIA – radioimmunoassay.

### 3.4 Covariate analysis

We next aimed to evaluate the influence of different covariates (year, sex and age) on FGM concentrations using the three immunoassays (**Figure 3A-F**). Our findings reveal that both the RIA and ELISA from AA were able to detect yearly variations in FGM concentrations as expected with a marked decrease in 2019 (**Figure 3D-E**), while the ELISA from CCC did not (**Figure 3F**). This suggests that the RIA and ELISA from AA are better suited for quantifying yearly stress variations in FGM extracts from yellow-bellied marmots.

**FIGURE 3.**
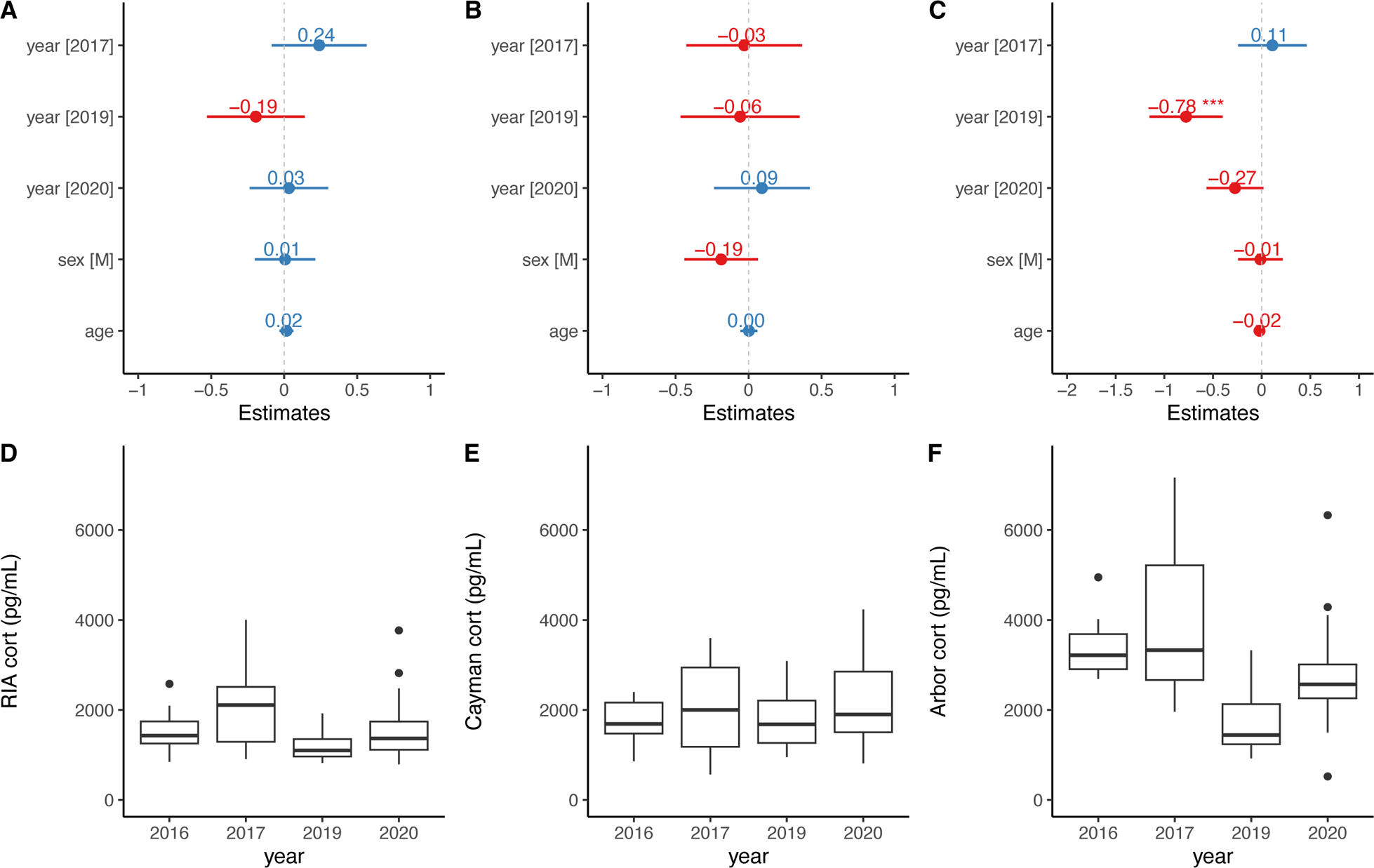
Evaluation of year, sex and age as covariates of corticosterone concentrations from yellow-bellied marmots’ fecal glucocorticoid metabolites (FGM) using a radioimmunoassay (RIA) **(A, D)**, and a competitive enzyme-linked immunosorbent assay (ELISA) from Arbor Assays **(B, E)** or Cayman Chemical Company **(C, F).**

### 3.5 Analytical validation of Arbor Assay ELISA

Because the AA ELISA had the best assay parameters as well as a close approximation to the RIA, we next tested its suitability with marmot feces using spike recovery, dilution linearity and parallelism experiments (Taha, 2024) using two different fecal samples (undiluted concentration = 662.8 ± 0.0 and 381.1 ± 89.0 pg/mL). Most spike recovery (**Figure 4A**), dilution linearity (**Figure 4B**) and parallelism (**Figure 4C**) experiments showed acceptable recoveries within the ± 20.0% range. Furthermore, we observed strong relative accuracy between the two diluted samples and the standard calibrator (**Figure 4D**). Collectively, these results indicate strong suitability and lack of matrix effects between the AA ELISA and yellow-bellied marmot feces.

**FIGURE 4.**
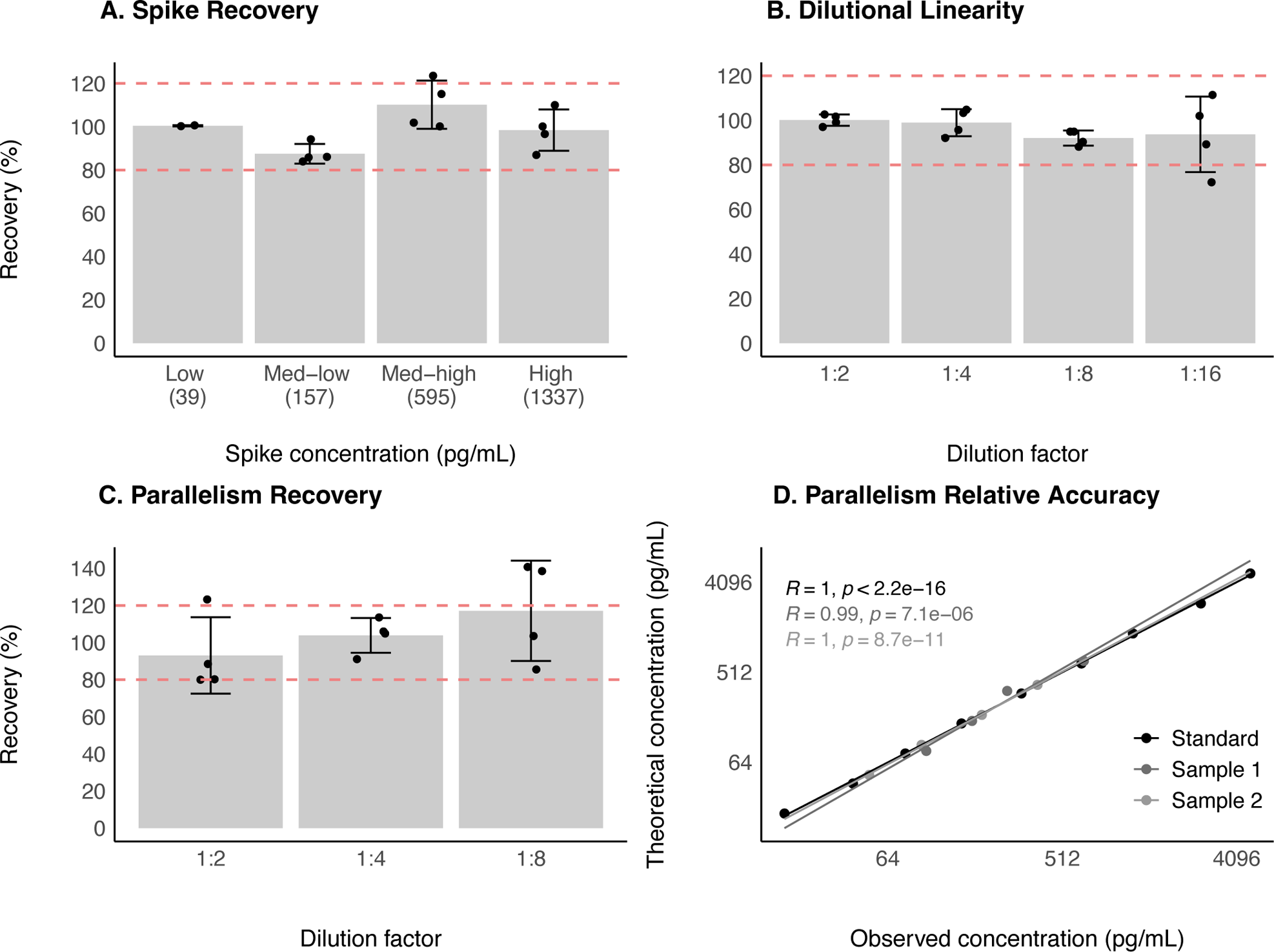
Analytical validation of the competitive enzyme-linked immunosorbent assay (ELISA) from Arbor Assays for quantification of corticosterone in yellow-bellied marmots’ fecal glucocorticoid metabolites extracts using spike recovery **(A)**, dilutional linearity **(B)** and parallelism **(C, D).**

## 4. DISCUSSION

The precise measurement of physiological stress levels in yellow-bellied marmots is critical, not only for understanding individual and population health but also for gaining insights into broader ecological dynamics (Dantzer et al., 2016; Kroeger et al., 2021; Pinho et al., 2019; Price et al., 2018). These marmots are particularly sensitive to environmental stressors, which can have profound effects on their survival and reproduction (Blumstein et al., 2016). Traditionally, RIAs, despite being less readily available nowadays, have been used extensively due to their early development and initial sensitivity for measuring FGMs including corticosterone (Hare et al., 2014; Keay et al., 2006; von der Ohe et al., 2004; Wasser et al., 2000). For over two decades, we relied on RIAs to measure corticosterone as a proxy for stress levels using yellow-bellied marmots’ FGM extracts, but unforeseen circumstances have compelled us to discontinue its use. Moreover, RIAs involve the use of radioactive materials, which pose safety concerns and are increasingly difficult to access due to stricter regulations and the declining availability of necessary isotopes. They also require elaborate safety measures and disposal procedures that complicate their use.

To ensure a continuity and comparability with decades of data while addressing the need for a safer, more sustainable method, we evaluated the use of two ELISAs in measuring corticosterone from yellow-bellied marmots’ FGM extracts, for which no analytically validated immunoassay is currently readily available. Comparison of corticosterone levels revealed that while the CCC ELISA often exceeded the highest standard and could not reliably quantify high concentrations, both the RIA and AA ELISA provided consistent results across the range, with the AA ELISA showing superior consistency and reliability (**Figure 1**). In analyzing immunoassay parameters, the AA ELISA outperformed the CCC ELISA with higher sensitivity, lower variability (**Table 1**) and error (**Tables 2-3**). Correlation analysis revealed strong associations between the AA ELISA and the RIA, suggesting the AA ELISA as the most accurate alternative to the RIA (**Figure 2A-C**). Finally, our covariate analysis highlighted the ability of the AA ELISA to detect expected yearly variations in FGM levels, further establishing its utility for long-term monitoring of stress in these marmots (**Figure 3D-F**). Given the positive outcomes and the close association with the RIA results, we proceeded with further analytical validation of the AA ELISA, specifically examining it for matrix effects. We found none, affirming the AA ELISA’s robustness and accuracy for use in measuring physiological stress levels in yellow-bellied marmots’ FGM extracts.

It is important to note that when comparing the concentrations of corticosterone in FGMs from all three immunoassays described here alongside other published studies, there was a lack of consensus on the true FGM concentration of the samples. For example, in our hands, the RIA, CCC and AA ELISAs yielded a concentration between 792–4004, 566–4234 and 522.8–7166 pg/mL, respectively. On the other hand, Price et al. (Price et al., 2018) used a similar AA ELISA and reported values between 5700–6900 pg/mL (converted from ng/g assuming a density of 1g/mL for FGMs). Moreover, Mateo et al. (Mateo and Cavigelli, 2005) quantified corticosterone from FGMs in captive Belding’s ground squirrels (*Spermophilus beldingi),* a species within the Sciuruinae subfamily, similarly to yellow-bellied marmots using RIA. They reported much lower values as depicted in their Figure 4B (range not provided). Moreover, whereas the CCC ELISA did not work well with our samples, other studies have documented its suitability with their samples. For instance, Holding et al. (Holding et al., 2020) showed that the CCC ELISA was suitable for quantifying stress levels in FGMs isolated from California ground squirrels (*Otospermophilus beecheyi*). A possible reason for this discrepancy could be due to different methodologies between our group and theirs in isolating FGMs.

## 5. CONCLUSIONS

We conclude that the AA ELISA is an appropriate kit to use when evaluating FGMs in wild yellow-bellied marmots. The immunoassay demonstrated strong association with the RIA kit in accurately detecting FGMs across sex and age, even in samples that had been extracted 7 years previously. We also analytically validated this immunoassay, showcasing the lack of matrix effects with the AA ELISA. Investigators interested in switching to a safer non-radioactive and easier to handle immunoassays for quantifying stress levels in corticosterone are encouraged to use the AA ELISA. Moreover, while demonstrating that long-term datasets can switch from RIA to ELISA, we urge future developments to keep longitudinal datasets in mind, maintain access and support for older immunoassays, and clearly communicate any changes.

## AUTHOR CONTRIBUTIONS

HBT, XOR and DTB conceptualized and designed the study. XOR curated the data, managed the project, and created the figures. XOR, EP, and HBT performed the experiments and collected the data. XOR and HBT analyzed the data. HBT, XOR and SR wrote the first draft of the manuscript. DTB provided data, funding, discussed results, edited and approved the final manuscript draft.

## ACKNOWLEDGEMENTS

We thank all the marmoteers who collected these samples over the years as well as the assay companies for providing guidance on using their kits.

## DATA SHARING

All data and code has been posted in a public OSF repository at https://osf.io/umqvn/ DOI: 10.17605/OSF.IO/UMQVN

